# A FRET-Based FLIM Method to Probe Membrane-Induced Alpha-Synuclein Aggregation in Neurons

**DOI:** 10.1101/2024.12.19.629536

**Authors:** Paula-Marie E. Ivey, Abdelrahman Salem, Sehong Min, Wenzhu Qi, Magaly Guzman Sosa, Tamara L. Kinzer-Ursem, Jean-Christophe Rochet, Kevin J. Webb

## Abstract

Parkinson’s disease (PD) involves the aggregation of the protein alpha-synuclein, a process promoted by interactions with intracellular membranes. To study this phenomenon in neurons for the first time, we developed a fluorescence lifetime imaging (FLIM) method using Förster resonance energy transfer and self-quenching reporters, analyzed with a custom-built FLIM microscope. This method offers insights into aggregate formation in PD and can be broadly applied to probe protein-membrane interactions in neurons.

## 2 Results

Parkinson’s disease (PD) is a neurodegenerative disorder characterized by the aggregation of the presynaptic protein alpha-synuclein (aSyn) in the brains of patients. Although aSyn aggregation has been extensively analyzed in recombinant protein solution, mechanisms underlying the initial self-assembly steps in neurons remain poorly understood. A substantial fraction of aSyn in neurons is bound to phospholipid membranes in an alpha-helical conformation [1, 2], where binding affinity is determined in part by the degree of membrane curvature [3, 4]. Additionally, certain orientations of membrane-bound aSyn can lead to the protein’s central hydrophobic region being exposed to the aqueous milieu, where it is available for interactions with other aSyn subunits that in turn lead to aggregate formation on the membrane surface [5–9]. Therefore, intracellular membranes may act as a platform for aggregation to occur [4, 10]. Prior work has demonstrated that aSyn can form aggregates on artificial membrane surfaces [7–9, 11, 12]. However, this mechanism has yet to be demonstrated in a cellular system, where multiple membranes can act as potential sites for aSyn binding. Here, we investigated aSyn aggregation in relation to the protein’s binding to early and late endosomes in cultured neurons.

We developed an imaging method by combining FRET- and self-quenching-induced lifetime changes to probe the role of lipid membranes in facilitating aSyn aggregation in neurons. Figure 1 (a) illustrates the FRET approach used to monitor the binding of aSyn to intracellular membranes. The method involves transducing primary rat cortical neurons with viruses encoding (i) a ‘marker’ protein localized to the lipid membrane of interest fused to mTurquoise2 (mT2) (FRET donor), and (ii) aSyn fused to mVenus (FRET acceptor). aSyn binding is detected via a decrease in mT2 lifetime. Figure 1 (b) shows the self-quenching phenomenon used to monitor aSyn-mVenus aggregation via fluorescence lifetime measurements [13]. We implemented this combined FRET and self-quenching imaging scheme to study aSyn-membrane inter-actions and membrane-induced aSyn aggregation in relation to early and late endosomes. The marker proteins selected for labeling the early and late endosomes were Rab5 and Rab7, respectively [2, 14]. Additionally, we used an aSyn-mVenus construct with the A53T mutation because this variant has a high aggregation propensity. To carry out the FLIM measurements, a custom time-gated wide-field FLIM system was constructed around an Olympus microscope platform (described in the Methods section).

**Fig. 1.**
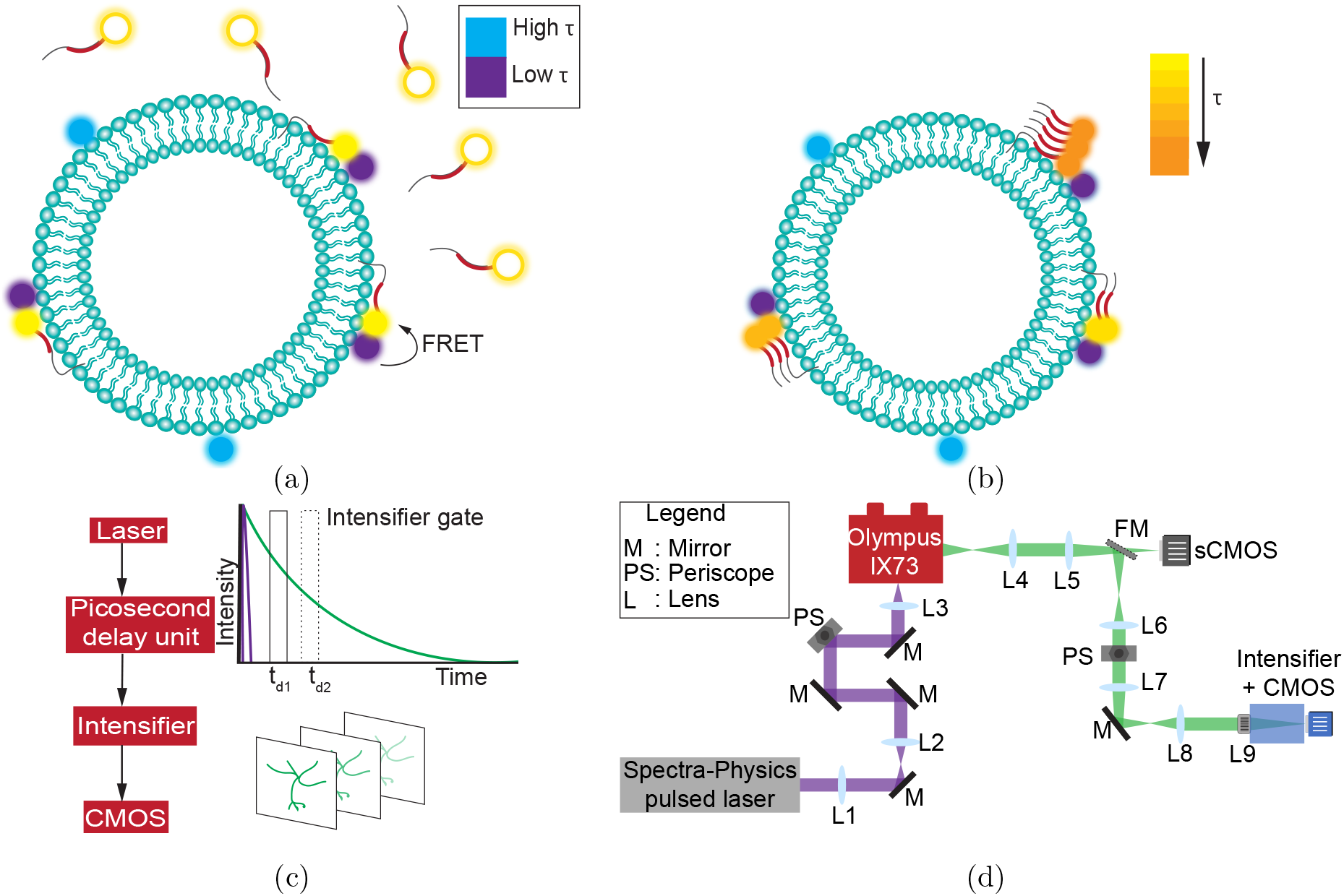
Combined FRET and self-quenching scheme for detecting aSyn membrane binding and aggregation. (a) FRET component, where the donor fluorophore lifetime (*τ*) decreases (light blue to dark blue) due to FRET between the membrane protein-mT2 and aSyn-mVenus. Upon exciting the donor, an mVenus acceptor located within the FRET distance becomes excited and can emit fluorescence (empty and solid yellow circles indicate non-fluorescent and donor-excited aSyn-mVenus, respectively). (b) Self-quenching component, where a decrease in aSyn-mVenus lifetime reports on the degree of aggregation due to a local concentration increase of the fluorophore. (c) Diagram of the lifetime measurement principle showing that an intensifier gate incrementally samples the fluorescence decay curve (green), and a picosecond delay unit changes the temporal gate delay relative to the laser excitation pulse (purple). At each gate delay position, an image is generated. (d) Schematic of the time-domain FLIM system used for detecting aggregation (top view, not drawn to scale). A Spectra-Physics laser coupled to an optical parametric amplifier system generates excitation light (purple), which is then directed to an Olympus iX73 microscope via a dichroic mirror. The path of emission light collected from the microscope is depicted in green. L: lens, M: mirror, BS: beam splitter, PS: periscope, FM: flip mirror.

To investigate interactions between aSyn and early endosomes and their impact on the protein’s aggregation, cortical neurons were transduced with adenovirus encoding Rab5-mT2 and aSyn-mVenus. A subset of cultures were also exposed to aSyn preformed fibrils (PFFs) to assess the impact of seeding on the aggregation of aSyn bound to early endosomes. As a FRET control, additional cultures were transduced with Rab5-mT2 virus alone, enabling the expression of the donor fluorophore without the aSyn-mVenus acceptor. After 5 days, the cells were fixed and analyzed via FLIM. The aSyn-mVenus lifetime was lower in neurons cultured in the presence versus the absence of PFFs, confirming the stimulatory effect of PFFs on intracellular aSyn aggregation (Figure 2(a and d)). In PFF-treated neurons, the Rab5-mT2 and aSyn-mVenus fluorescence signals showed overlapping spatial distributions. However, the Rab5-mT2 lifetime in PFF-treated neurons was similar to that in neurons cultured without PFFs, which did not exhibit a clear overlap in mVenus and mT2 fluorescence (Figure 2(a)). Furthermore, no differences in Rab5-mT2 lifetime were observed in neurons co-expressing Rab5-mT2 and aSyn-mVenus, whether cultured with or without PFFs, compared to control neurons expressing Rab5-mT2 alone (Fig. 2(b and c)). These results suggest that FRET does not occur between Rab5-mT2 and aSyn-mVenus, indicating the absence of a strong interaction between aSyn and early endosomes in this neuronal model, regardless of the presence of aSyn seeds.

**Fig. 2.**
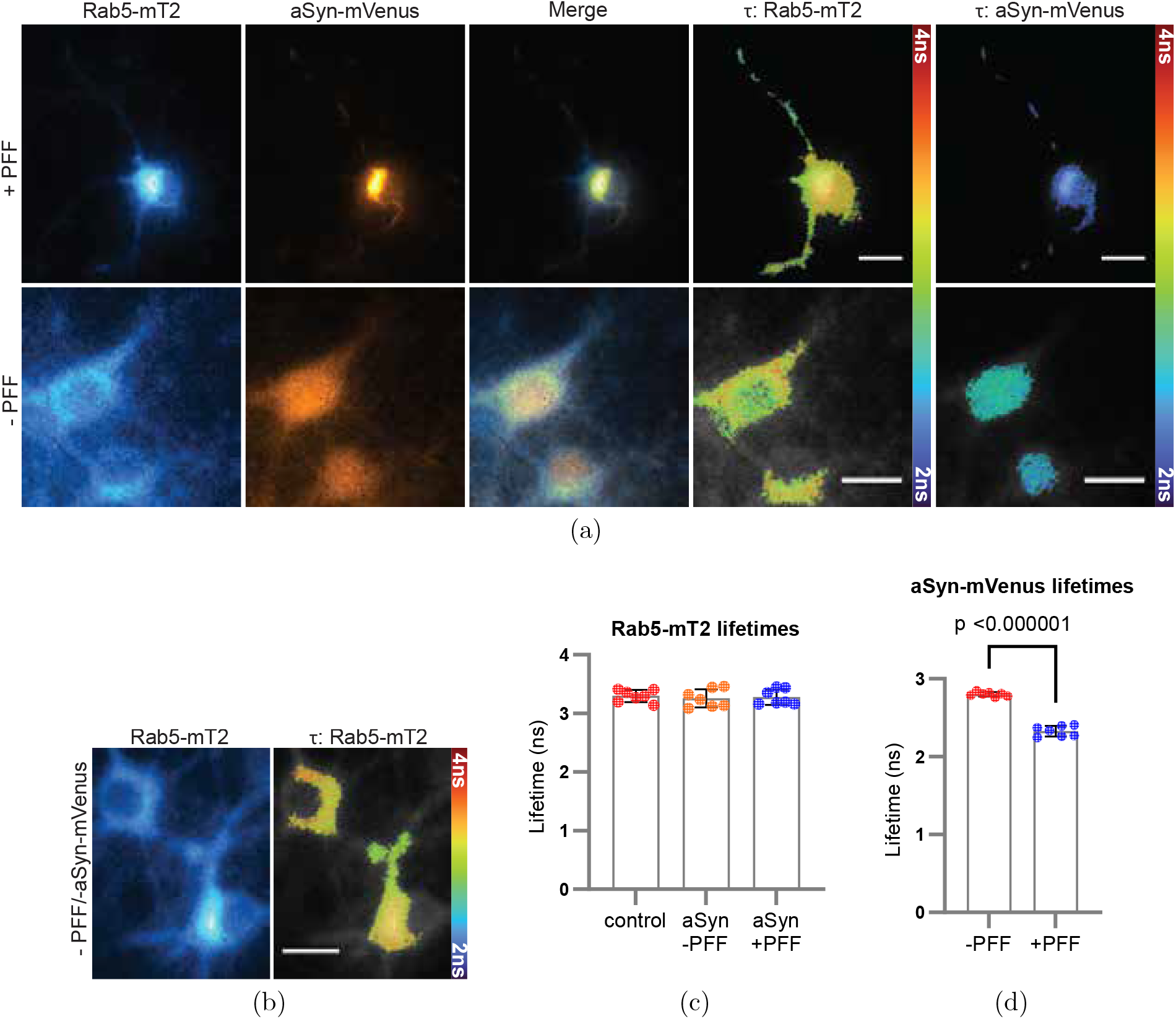
FLIM-FRET analysis indicates no interaction between aSyn-mVenus and Rab5-mT2 in neurons, regardless of aSyn PFF treatment. (a) Images of fixed cortical neurons transduced with adenoviruses encoding Rab5-mT2 and aSyn-mVenus in the presence (top row) or absence (bottom row) of aSyn PFFs, recorded 5 days post-PFF treatment. The images display (from left to right) the fluorescence intensity of Rab5-mT2 and aSyn-mVenus, the merged intensity signals, and the corresponding Rab5-mT2 and aSyn-mVenus lifetime maps. (b) Images of fixed cortical neurons transduced with adenovirus encoding Rab5-mT2 in the absence of aSyn PFFs, showing Rab5-mT2 fluorescence intensity (left) and the corresponding lifetime map (right). Scale bars in (a) and (b): 10 *µ*m. (c) Graph showing the Rab5-mT2 fluorescence lifetime for neurons expressing Rab5-mT2 alone (‘control’) or Rab5-mT2 and aSyn-mVenus, without or with aSyn PFF treatment. (d) Graph showing the aSyn-mVenus fluorescence lifetime for neurons co-expressing Rab5-mT2 and aSyn-mVenus, without or with aSyn PFF treatment. The data in (c) and (d) are plotted as the mean +*/*− SEM, *n* = 7 neurons per group. Statistical analyses consisted of ANOVA in (c), showing no significant differences among means, and an unpaired t-test in (d).

Next, we examined interactions between aSyn and late endosomes, along with their influence on aSyn aggregation, using the same experimental setup and imaging approach as described for early endosomes. The aSyn-mVenus lifetime was lower in neurons cultured in the presence versus the absence of PFFs, confirming seeded aggregation of the protein in response to the PFF treatment (Figure 3(a and d)). PFF-exposed neurons showed extensive overlap of the Rab7-mT2 and aSyn-mVenus fluorescence in both the soma and neurites. Additionally, the Rab7-mT2 lifetime showed a trend towards being reduced in PFF-treated neurons, especially in neuritic regions, compared to neurons cultured without PFFs, where the overlap of mVenus and mT2 fluorescence was less pronounced (Figure 3(a and c)). Even in the absence of PFFs, aSyn-mVenus-expressing neurons exhibited a decreased Rab7-mT2 lifetime compared to control neurons expressing Rab7-mT2 alone (Fig. 3(b and c)). These observations suggest that FRET occurs between Rab7-mT2 and aSyn-mVenus, in turn implying that aSyn interacts to a greater extent with late versus early endosomes in this neuronal model, particularly when aSyn seeds are present.

**Fig. 3.**
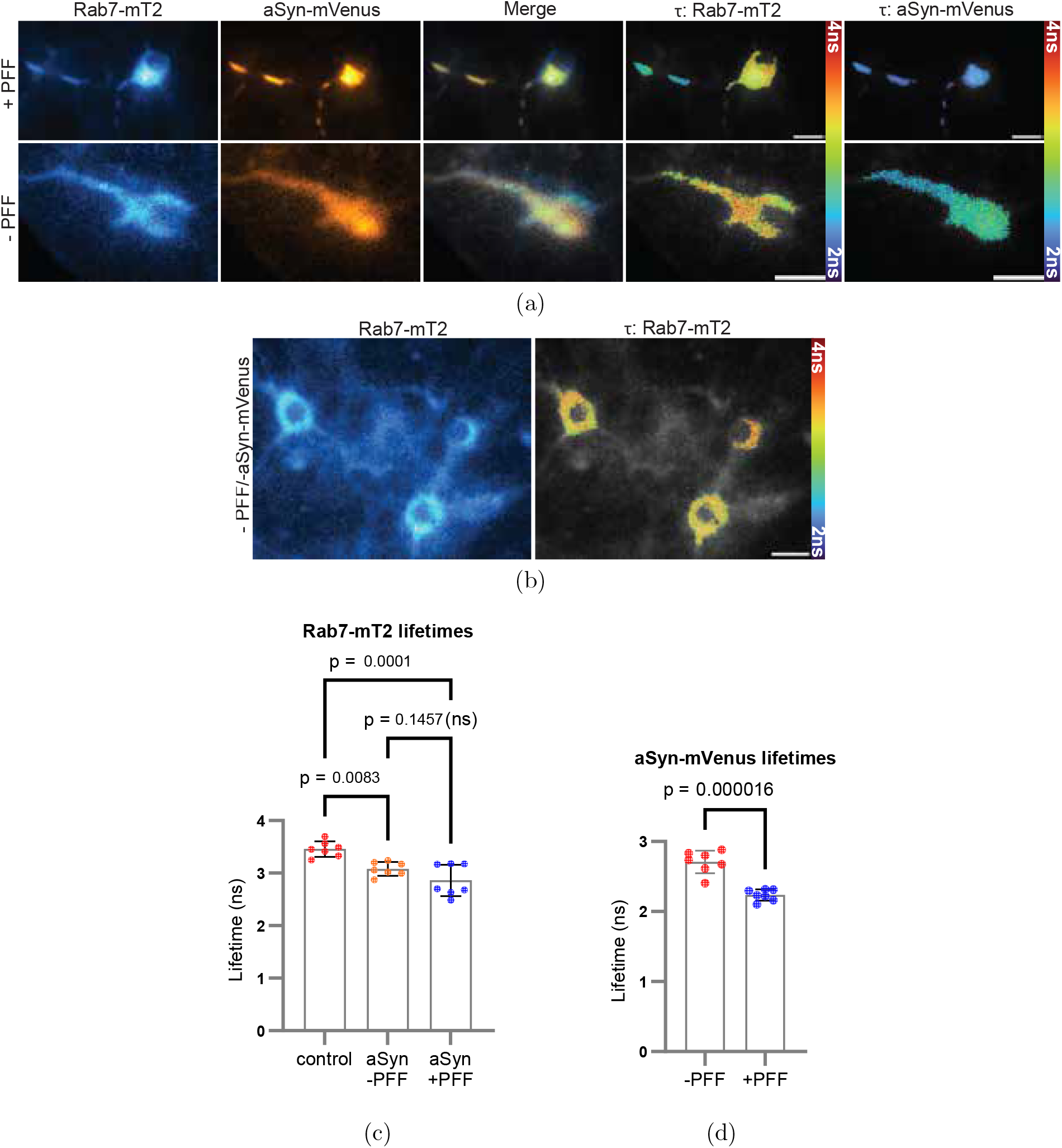
FLIM-FRET analysis indicates an interaction between aSyn-mVenus and Rab7-mT2 in neurons cultured with or without aSyn PFFs. (a) Images of fixed cortical neurons transduced with adenoviruses encoding Rab7-mT2 and aSyn-mVenus in the presence (top row) or absence (bottom row) of aSyn PFFs, recorded 5 days post-PFF treatment. The images display (from left to right) the fluorescence intensity of Rab7-mT2 and aSyn-mVenus, the merged intensity signals, and the corresponding Rab7-mT2 and aSyn-mVenus lifetime maps. (b) Images of fixed cortical neurons transduced with adenovirus encoding Rab7-mT2 in the absence of aSyn PFFs, showing Rab7-mT2 fluorescence intensity (left) and the corresponding lifetime map (right). Scale bars in (a) and (b): 10 *µ*m. (c) Graph showing the Rab7-mT2 fluorescence lifetime for neurons expressing Rab7-mT2 alone (‘control’) or Rab7-mT2 and aSyn-mVenus, without or with aSyn PFF treatment. (d) Graph showing the aSyn-mVenus fluorescence lifetime for neurons co-expressing Rab7-mT2 and aSyn-mVenus, without or with aSyn PFF treatment. The data in (c) and (d) are plotted as the mean +*/*− SEM, *n* = 7 neurons per group. Statistical analyses consisted of ANOVA with Tukey’s post hoc test in (c) and an unpaired t-test in (d).

Experiments with neurons co-expressing Rab5- or Rab7-mT2 and mVenus (not fused to aSyn) in the absence of aSyn PFFs demonstrated that the mT2 lifetime remained unchanged compared to that measured in control neurons expressing Rab5- or Rab7-mT2 alone (Supplementary Fig. 1). Conversely, the mT2 lifetime was slightly reduced in PFF-treated neurons co-expressing Rab5- or Rab7-mT2 and mVenus, compared to control neurons expressing Rab5- or Rab7-mT2 without PFF treatment. These findings suggest that aSyn-mVenus binding to late (Rab7-positive) endosomes (Fig. 3 (c)) is not driven primarily by the mVenus module. Although non-specific mVenus-membrane interactions may have contributed to the observed PFF-dependent binding of aSyn-mVenus to late endosomes, our observation that mVenus also bound to early (Rab5-positive) endosomes in PFF-treated neurons (Supplementary Fig. 1 (a)), in contrast to aSyn-mVenus (Fig. 2 (c)), suggests that mVenus may not adopt the same fold as an isolated protein compared to when it is fused to aSyn. Lastly, the mVenus lifetime was unchanged in neurons cultured in the presence versus the absence of PFFs (Supplementary Fig. 2), indicating that the reduced aSyn-mVenus lifetime observed in PFF-treated neurons (Figs. 2 (d) and 3 (d)) was attributable to aSyn aggregation.

Our results show that A53T aSyn associates preferentially with late endosomes versus early endosomes. This association shows a trend towards being increased in neurons treated with aSyn seeds, suggesting that late endosomes may act as a platform for seeded aggregation. Overall, we demonstrate the benefit of using a joint self-quenching and FRET lifetime imaging method to probe protein-membrane interactions in relation to protein self-assembly in neurons.

## 3 Methods

### 3.1 Fluorescence Lifetime Imaging Microscopy

A custom time-domain FLIM setup was constructed around an Olympus iX73 inverted microscope platform. The excitation source was a 1-MHz 1040-nm pump laser coupled with an optical parametric amplifier (Spectra-Physics) to support a broader range of excitation wavelengths (320-1040 nm). The laser was coupled to the microscope with a series of lenses and mirrors. The system was equipped with two image-capturing devices, an sCMOS camera (Photometrics PRIME) capable of up to 100 fps collection rates for intensity-based imaging, and a high-rate image-intensified (PicoStar HR12, LaVision) CMOS (Imager-M-Lite, LaVision) integrated system for fluorescence lifetime recording. A picosecond delay unit (PSD) was used to introduce delays between the intensifier gate and the exciting laser pulse. The delay unit used the electrical trigger output of the laser. For the FLIM measurements described below, an intensifier gate width of 200 ps was applied and delayed relative to the laser pulse in 200 ps increments to sample the fluorescence decay curve. A pixel-wise mono-exponential fit with the deconvolved temporal gate function was then performed on the data to determine the lifetime.

### 3.2 Expression and Purification of Recombinant aSyn Protein

Recombinant aSyn was purified as previously described [15, 16]. *E. coli* BL21 (DE3) cells were transformed with the bacterial expression vector pT7-7 encoding mouse aSyn. A single colony was used to inoculate LB medium supplemented with ampicillin (100 *µ*g/L), and the culture was incubated at 37°C until the optical density at 600 nm reached 0.5-0.6. Protein expression was induced with isopropyl *β*-D-1-thiogalactopyranoside at a final concentration of 1 mM, followed by incubation at 37°C for 4 h. The cells were then harvested by centrifugation, resuspended in lysis buffer A (10 mM Tris HCl, pH 8.0, 1 mM EDTA, 0.25 mg/mL lysozyme), and lysed using a French press cell disruptor (Thermo Electron, Waltham, MA) at *>* 1000 psi. The lysate was treated with 0.1% (w/v) streptomycin sulfate to precipitate DNA and then clarified by centrifugation. The supernatant was subjected to partial purification via two successive ammonium sulfate precipitations (30% and 50% saturation) at 4°C. The resulting pellet was resuspended in 10 mM Tris-HCl (pH 7.4), and the suspension was heated to 100°C for 15 min. Following centrifugation at 13,500 × g at 4°C for 20 min to precipitate denatured proteins, the supernatant was filtered through a 0.22-*µ*m membrane. aSyn was purified via successive fractionations using (i) a HiLoad 16/600 Superdex 200 pg size exclusion column (Cytiva, Marlborough, MA), with elution performed in 10 mM Tris HCl (pH 7.4); and (ii) a HiPrep Q HP 16/10 anion-exchange column (Cytiva), with elution performed using a linear gradient of 25 mM to 1 M NaCl in a buffer consisting of 10 mM Tris HCl (pH 7.4) and 1 mM EDTA. Fractions enriched with aSyn (identified via SDS-PAGE with Coomassie blue staining) were pooled, and the solution was dialyzed against PBS (10 mM phosphate buffer, 2.7 mM KCl, and 137 mM NaCl, pH 7.4). The purified protein was stored at −80°C until use, with a final purity of approximately 95%.

### 3.3 Preparation of aSyn Preformed Fibrils

Purified mouse aSyn was concentrated to 5 mg/mL (347 *µ*M) using a 10 kDa molecular weight cutoff (MWCO) spin filter (Sartorius, Columbus, OH) operated at 4,500 × g. The concentrated protein solution (0.5 mL) was filtered through a 0.22-*µ*m syringe filter (Fisherbrand polyethersulfone, 33 mm) and incubated in a sterile 1.5-mL microcentrifuge tube at 37°C for 7 days with continuous shaking at 123 × g in a Thermomixer (BT LabSystems BT917). The resulting fibrils were collected by centrifugation at 13,000 × g for 15 min and resuspended in 250 *µ*L of Dulbecco’s phosphate-buffered saline (DPBS; Cytiva). An aliquot of the fibril suspension was treated with guanidine hydrochloride (final concentration, 8 M) and incubated at 22°C for 1 h to dissociate fibrils into monomers. The aSyn concentration was measured via absorbance at 280 nm using a Nanodrop spectrophotometer, with an extinction coefficient of 7450 M^-1^cm^-1^. Fibrils were resuspended to a final concentration of 5 mg/mL, divided into 257 *µ*L aliquots, and stored at −80°C. Before use, fibril suspensions were sonicated in ethanol-sterilized tubes (Active Motif) using a cup horn sonicator (Qsonica Q700) set to 30% power (100 W/s) with a cycle of 3 s on and 2 s off, for a total ‘on’ time of 8 min. Fibrils underwent a total of 2 rounds of sonication. The bath temperature was maintained between 5°C and 15°C throughout the sonication.

### 3.4 Preparation of Adenoviral Constructs

Adenoviral constructs encoding aSyn A53T-mVenus, Rab5-mT2, Rab7-mT2, or mVenus were produced using the Virapower Adenoviral Expression System from Invitrogen (Carlsbad, CA). cDNAs encoding mVenus or mT2 were generated via PCR using the plasmids ecAT3.10 (Addgene #107215) and pRSETB-tdTomato-*ε*-CeN [17], both provided by Dr. Mathew Tantama (Wellesley College), as templates. cDNAs encoding human aSyn, Rab5b, and Rab7a were prepared via PCR using the plasmids pENTR-aSyn [7], mCh-Rab5 (Addgene #49201, deposited by Dr. Gia Voeltz), and GFP-Rab7 WT (Addgene #12605, deposited by Dr. Richard Pagano) as templates. mCh-Rab5 and GFP-Rab7 WT were provided by Dr. Robert Stahelin (Purdue University). PCR reactions were carried out using Phusion High-Fidelity DNA polymerase or Q5 Hot Start High-Fidelity DNA polymerase (NEB, Ipswich, MA), and the sequences of the oligonucleotide primers used to generate the constructs are listed in the supplementary material (Supplementary Table 1). Overlapping PCR fragments encoding aSyn-mVenus, Rab5-mT2, or Rab7-mT2, and a single PCR fragment encoding mVenus, were subcloned into a variant of the entry vector pENTR1A carrying the human synapsin promoter (h-syn-P) [7], digested with KpnI and XhoI. The ligation reactions were carried out using the NEBuilder HiFi DNA Assembly kit (NEB). The pENTR1A-h-syn-P-aSynA53T-mVenus plasmid was generated by site-directed mutagenesis using overlap extension PCR to introduce the A53T mutation. The insert from each pENTR1A construct was transferred into the ‘promoter-less’ pAd/PL-DEST5 adenoviral expression vector [7] via Gateway recombination cloning (Invitrogen). The DNA sequence of each insert was verified by Sanger sequencing (Genewiz, South Plainfield, NJ). The resulting adenoviral constructs were packaged into virus via lipid-mediated transient transfection of the HEK 293A packaging cell line. Adenoviral titers were determined using the Adeno-X qPCR titration kit from Takara Bio USA (Mountain View, CA).

### 3.5 Preparation and Treatment of Primary Cortical Cultures

Primary cortical cultures were prepared as described [16] by dissecting day 17 embryos obtained from pregnant Sprague–Dawley rats (Envigo, Indianapolis, IN) using methods approved by the Purdue Animal Care and Use Committee. Briefly, the cortical layers were isolated stereoscopically and dissociated by incubation with papain (20 U/mL) in sterile Hank’s Balanced Salt Solution (HBSS) at 37°C for 45 min. The dissociated cells were plated on poly-D-lysine-coated 8-well chambered glass plates (Cellvis) at a density of 75,000 cells/well in Neurobasal media supplemented with 2% (v/v) B-27 supplement, 5% (v/v) FBS, 1% (v/v) GlutaMAX, 50 U/mL penicillin, and 50 *µ*g/mL streptomycin. The next day, the plating media was replaced with Neurobasal media plus 2% (v/v) B-27 supplement, 1% (v/v) GlutaMAX, 10 U/mL penicillin, and 10 *µ*g/mL streptomycin. After 6 days in vitro (DIV = 6), the cultures were transduced with adenovirus encoding aSyn A53T-mVenus, Rab5-mT2, Rab7-mT2, and/or mVenus under the control of the synapsin promoter at a multiplicity of infection (MOI) of 5. A subset of cultures were also treated with aSyn PFFs (final concentration, 6 *µ*g/mL). The cells were then incubated for 5 days, fixed with 4% (w/v) paraformaldehyde (PFA) in PBS for 15 min, and imaged via FLIM. mT2 and mVenus fluorescence lifetime data were recorded from 5 fields of view per well of the 8-well chambered glass plate at a magnification of 20X or 40X. mT2 was excited at 430 nm, with emission collected through a 497/20 nm band-pass filter. mVenus was excited at 488 nm, with emission collected through a 530/40 nm band-pass filter.

### 3.6 Statistical Analysis

The data were analyzed with an unpaired t-test (for experiments with 2 groups) or a one-way ANOVA with Tukey’s multiple comparisons post hoc test (for experiments with *>* 2 groups) using GraphPad Prism version 8.0 (La Jolla, CA).

## Acknowledgments

This work was supported in part by the National Science Foundation (CBET 1937986 and EAGER 2330643), the National Institutes of Health (R21 NS105048 and NS135424), and the Michael J. Fox Foundation.

## Declarations

- Funding: National Science Foundation (CBET 1937986 and EAGER 2330643), the National Institutes of Health (R21 NS105048 and NS135424), and by the Michael J. Fox Foundation.
- Competing interests: The authors declare no competing interests
- Data availability: Data available upon request
- Author contribution: P.M.E.I conceived and designed the study with guidance from J.C.R and K.J.W.P.M.E.I designed and built the FLIM system and carried out the experiments. A.S. developed the algorithms for imaging and data analysis. A.S and P.M.E.I analyzed and interpreted the data. S.M. developed and prepared the viral constructs. W.Q. prepared the neuron samples for the studies. M.G.S prepared the PFFs. P.M.E.I wrote the manuscript with guidance from J.C.R, K.J.W, and T.L.K.U.

