## Supplementary Material for "A FRET-Based FLIM Method to Probe Membrane-Induced Alpha-Synuclein Aggregation in Neurons"

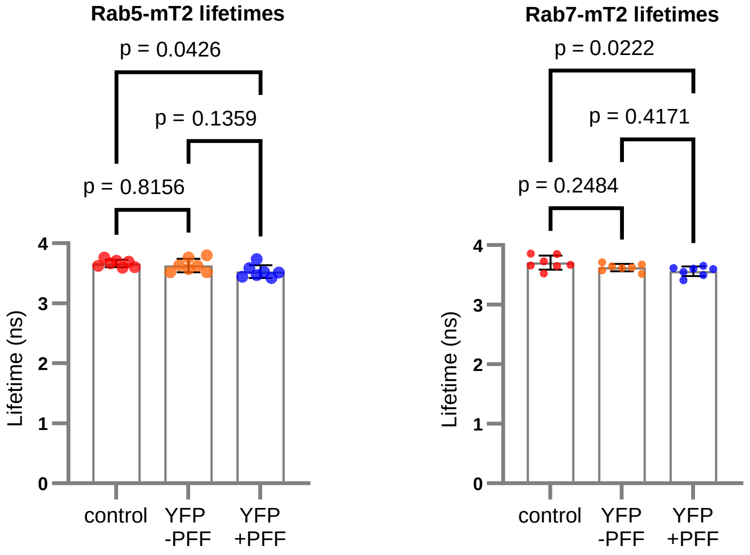


(a) (b)

**Fig. 1** FLIM-FRET analysis indicates no interaction between mVenus and Rab5-or Rab7-mT2 in neurons cultured without aSyn PFFs. (a) Graph showing the Rab5-mT2 fluorescence lifetime for neurons expressing Rab5-mT2 alone ('control') or Rab5-mT2 and mVenus ('YFP'), without or with aSyn PFF treatment. (b) Graph showing the Rab7-mT2 fluorescence lifetime for neurons expressing Rab7-mT2 alone ('control') or Rab7-mT2 and mVenus ('YFP'), without or with aSyn PFF treatment. The data in (a) and (b) are plotted as the mean +/− SEM, $n = 7$ neurons per group. Statistical analyses consisted of ANOVA with Tukey's post hoc test. A slight decrease in Rab5- or Rab7-mT2 fluorescence lifetime was observed in neurons co-expressing Rab5- or Rab7-mT2 and mVenus, in the presence but not the absence of aSyn PFFs, compared to control neurons expressing Rab5- or Rab7-mT2 without mVenus co-expression or PFF exposure.

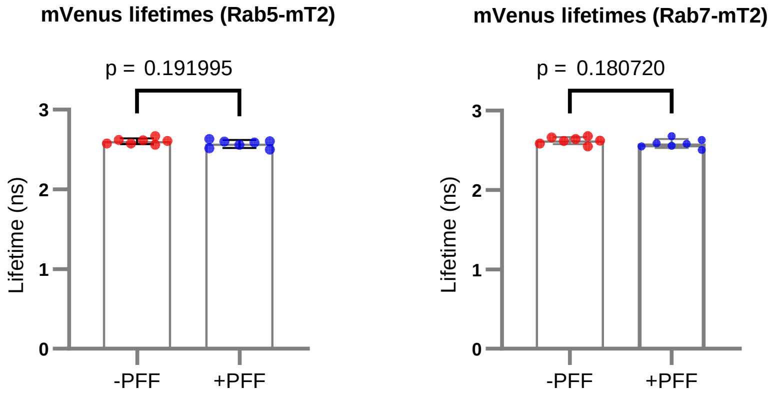


(a) (b)

**Fig. 2** FLIM-FRET analysis indicates no mVenus self-assembly in neurons incubated with aSyn PFFs. The graphs show the aSyn-mVenus fluorescence lifetime for neurons co-expressing Rab5-mT2 (a) or Rab7-mT2 (b) and mVenus, without or with aSyn PFF treatment. The data are plotted as the mean +/− SEM, $n = 7$ neurons per group. Statistical analyses consisted of unpaired t-tests.

| Primer | Sequence (5'-3') |
|---|---|
| aSyn-linker-mVenus-For | GACTACGAACCTGAAGCCGCTCCAGTAGCTACAATGGTGAGCAAGGGCG |
| mVenus-XhoI-pENTR-Rev | GCTGGGTCTAGATATCTCGAGTTACTTGTACAGCTCGTCCATG |
| pENTR-hsyn-KpnI-For | CAAGGATCCACCGGTACCGCCGCCAC |
| aSynA53T-Rev | CCACTGTTGTCACACCATGCACCACTCC |
| aSynA53T-For | GGTGTGACAACAGTGGCTGAGAAGACCAAAG |
| aSyn-EcoRI-Rev | CCACAGGCATATCTTCCAGAATTC |
| pENTR-hSyn-mTurquoise2-For | AGGATCCACCGGTACCGCCGCCACCATGGTGAGCAAGGGCGAGGAG |
| mTurquoise2-APVAT-Rev | TGTAGCTACTGGAGCAATCCCGGCGGCGGTCACGAA |
| APVAT-Rab5b-For | GCTCCAGTAGCTACAATGACTAGCAGAAGCACAGCTAG |
| Rab5b-pENTR-Rev | TTACTAACCGGTACGCTCGAGTCAGTTGCTACAACACTGGC |
| APVAT-Rab7-For | GCTCCAGTAGCTACAACCTCTAGGAAGAAAGTG |
| Rab7-pENTR-Rev | TTACTAACCGGTACGCTCGAGTCAGCAACTGCAGCTTT |
| mVenus-KpnI-pENTR-For | CCGGTACCGCCGCCACCATGGTGAGCAAGGGCG |

**Table 1** Sequences of the oligonucleotide primers used to generate the adenoviral constructs.
